# Single Cell Proteomics in the Developing Human Brain

**DOI:** 10.1101/2025.06.23.660079

**Authors:** Tianzhi Wu, Lihua Jiang, Tanzila Mukhtar, Li Wang, Ruiqi Jian, Cheng Wang, Tiffany Trinh, Arnold R. Kriegstein, Michael Snyder, Jingjing Li

**Author notes:** Correspondence should be addressed to: ARK, MS, JL. Contribute equally to this project (Tianzhi Wu, Lihua Jiang).

## Abstract

Proteins are the functional effectors of virtually all biological processes, and accurately measuring their abundance and dynamics is essential for understanding development and disease. Although mRNA levels have historically been used as proxies for protein expression, growing evidence, especially from studies of the human cerebral cortex, has revealed widespread discordance between transcript and protein abundance. To directly address this limitation, we developed a rigorously optimized workflow that combines single-cell mass spectrometry with precise sample preparation to resolve, for the first time, quantitative proteomes of individual cells from the developing human brain. Our platform achieved deep proteomic coverage (∼800 proteins per cell) even in immature prenatal human neurons (5–10 μm diameter, ∼100 pg of protein per cell), capturing major brain cell types and enabling proteome-wide characterization at single-cell resolution. This approach revealed extensive transcriptome–proteome discordance across cell types, with particularly strong discrepancies in genes associated with neurodevelopmental disorders, a finding validated through orthogonal experiments. Proteins exhibited markedly higher cell-type specificity than their mRNA counterparts, underscoring the importance of proteomic-level analysis for resolving cellular identity and function. By reconstructing developmental trajectories from radial glia to excitatory neurons at the proteomic level, we identified dynamic stage-specific protein co-expression modules and pinpointed the intermediate progenitor-to-neuron transition as a molecularly vulnerable phase linked to autism. Altogether, by enabling single cell proteomics, this study establishes a foundational resource and technological advance for developmental neuroscience. It demonstrates that single-cell proteomics can capture critical developmental events and disease mechanisms that are undetectable at the transcript level. As this technology continues to improve in sensitivity and scalability, single-cell proteomics will become an indispensable tool for uncovering the molecular logic of brain development and for illuminating pathophysiological processes underlying neurodevelopmental disorders.

## Introduction

Proteins are the cornerstone of life, orchestrating biological processes and executing the vital functions that sustain cellular activities. While genetic studies have advanced our understanding of complex diseases, translating these findings into mechanistic insights hinges on validation at the protein level, the ultimate functional unit of biology. Yet, mRNA expression has long served as a proxy for protein activity. Although mRNA provides a convenient snapshot of transcriptional activity, growing evidence reveals its limitations in accurately reflecting protein abundance^1,2^.

Our recent Genotype-Tissue Expression (GTEx) study^3^, analyzing RNA and protein levels across 201 samples from 32 human tissues, uncovered a median correlation coefficient of only 0.46, indicating that mRNA explains a mere ∼20% of variation in protein abundance, with the strongest discordance in the cerebral cortex. A subsequent Drosophila study echoed this observation, finding discordance in nearly every gene^4^. The implications of the mRNA-protein disconnect are profound. When protein-level responses diverge from mRNA patterns, transcriptomics-guided therapies may falter. Compounding this issue, the previously observed mRNA and protein discordance was mostly based on bulk tissues^5–8^, where cell-to-cell variability is averaged out. Consequently, the true scale of discordance is likely even more pronounced at the single-cell level, especially in dynamic systems like the rapidly developing brain. This study thus specifically targets the developing human brain to reveal its single-cell proteomics landscape. Building on single-cell epigenomic and transcriptomic atlases of the developing brain^9^, the single-cell proteomic dimension presented in this study bridges the chasm between genetic insights and functional biology, creating a unified framework to decode the molecular logic of brain development and its vulnerabilities.

Single-cell proteomics stands at a critical inflection point. Targeted approaches such as CyTOF^10^ and CITE-seq^11^ are widely used but inherently limited: they provide only semi-quantitative measurements, rely on antibody availability and specificity, and collectively capture less than 1% of the proteome. In contrast, untargeted mass spectrometry (MS) holds the theoretical potential to profile the entire proteome in a single run^12^. However, early implementations required large cellular inputs, detecting only ∼400 proteins per cell in exceptionally large cells such as ∼100 μm oocytes^13^. Subsequent innovations in multiplexing, using isobaric labeling and carrier channels, boosted protein coverage by pooling material from multiple cells^14^. Yet these gains came with trade-offs: ratio compression^15^, batch effects^16^, and increased data variability^17^, ultimately limiting the utility of these strategies for high-resolution tissue mapping. Recent advances in mass spectrometry hardware and acquisition strategies have enabled a new generation of label-free, direct single-cell proteomics^18^. These next-generation platforms can quantify hundreds to thousands of proteins from individual cultured cells such as HeLa cells (∼250 pg of protein), bypassing multiplex-specific artifacts and achieving impressive depth without sample pooling. Nevertheless, most applications to date remain confined to immortalized cell lines or modest-scale studies (∼100 cells)^19–21^, often focused on technical benchmarking rather than biological insight. A key open question is whether these emerging capabilities can scale to match the power and resolution of single-cell RNA sequencing in complex primary tissues. This challenge is especially acute in the developing human cortex, where fetal neurons measure just 5–10 μm in diameter and contain approximately 100 pg of total protein, well below the protein input levels validated in existing single-cell MS workflows. Given their diminutive size and low protein abundance, it remains unknown whether the current technological innovation can unlock new opportunities for mapping complex tissue architecture. Although recent commentaries express optimism that single-cell proteomics will “take center stage” in cellular and developmental biology^22^, its application to comprehensive tissue mapping remains largely unrealized^23^.

In this study, we combined a meticulously optimized experimental workflow with advanced single-cell mass spectrometry (MS) profiling to generate a high-resolution map of the single-cell proteome landscape in the developing human brain. This dataset distinguishes major cell types at protein resolution and exposes both concerted and divergent mRNA–protein relationships across cell types. Independent validation confirmed that the observed discordances reflect biological regulation, not technical bias. Strikingly, the proteins showing the greatest divergence from their mRNA profiles are enriched for genes implicated in neurodevelopmental disorders, underscoring the need for direct protein-level measurements when interrogating disease mechanisms. We further demonstrate that proteins exhibit sharper cell-type specificity than their corresponding mRNAs, that frequently exhibit broader and more diffuse expression patterns. Importantly, our single-cell proteomic dataset allowed us to reconstruct detailed developmental trajectories, tracking cellular transitions at the proteomic level from radial glia through intermediate progenitors to mature excitatory neurons. Within this developmental pathway, we discovered a tightly coordinated module of proteins whose abundance sharply increased starting at the intermediate progenitor stage. Notably, this protein module showed exceptional evolutionary conservation and strong association with autism spectrum disorders (ASD), highlighting a critical developmental window of vulnerability revealed by single-cell proteomic analysis. Collectively, this study presents the first single-cell proteomic data of human brain development, complementing the prevailing transcriptomics-focused paradigm in neurogenomic research. By bridging the gap between gene expression and functional protein states, our framework reveals the underlying regulatory logic of cell identity and uncovers disease-relevant molecular players that remain invisible to RNA-based approaches. This integrated proteogenomic strategy not only deepens our understanding of neurodevelopment but also sets a new paradigm for investigating complex diseases, with broad implications for biomarker discovery and therapeutic innovation.

## Results

We collected fresh prenatal human brain samples at gestational weeks (GW) 13, 15, and 19. For GW15 and GW19, we microdissected the germinal zone and cortical plate from the prefrontal cortex. For the GW13 sample, we used the entire nascent frontal telencephalon (including ganglionic eminence) (**Fig. 1A**). Following tissue dissociation, we used FACS to agnostically isolate individual cells into low protein-binding microwells, each preloaded with lysis buffer to ensure efficient single-cell lysis for downstream mass spectrometry (MS) profiling **(Materials and Methods).** We adopted a label-free strategy where we dedicated each MS run to a single cell, leveraging the enhanced single-cell sensitivity and resolution afforded by this technology. Protein abundances were quantified from MS2 fragment-ion intensities using DIA-NN^24^. In total, performing 2,310 independent MS runs, we resolved the quantitative proteome for each of 2,310 single cells.

**Fig 1.**
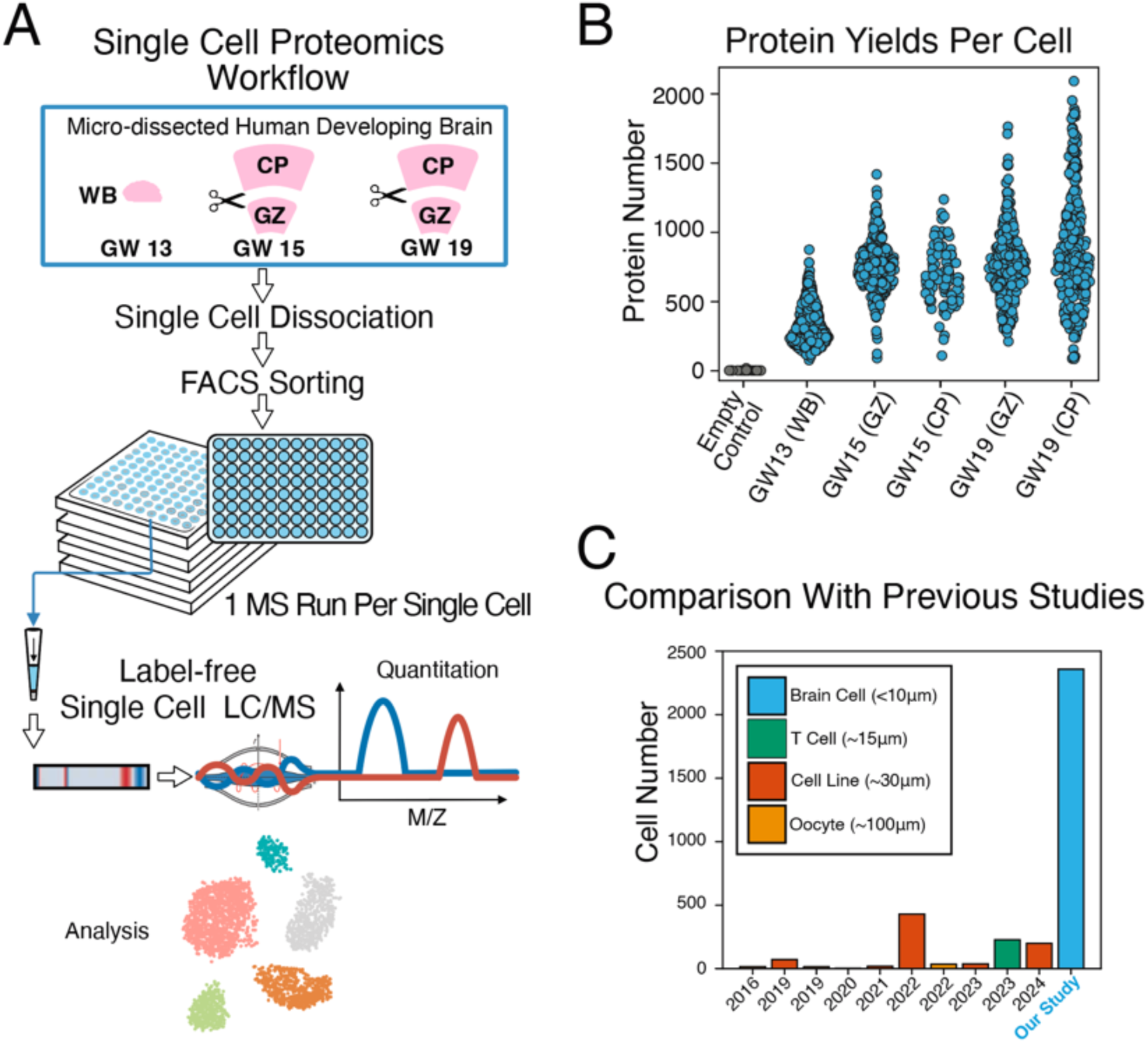
Label-free Single-Cell Proteome Profiling in Human Developing Brain Tissue. A. Label-free single-cell proteome workflow. Fresh tissue samples were immediately micro-dissected into GZ and CP region, or the whole frontal lobe in GW13 due to underdeveloped cortex. Individual cells were dissociated and sorted into low protein-binding microwells with preloaded buffer. Optimized label-free strategy was then applied to resolve the proteome for each single cell. **GW:** gestational week, **GZ:** germinal zone, **CP:** cortical plate, **WB**: whole brain tissue. B. Approximately 800 proteins per single cell were resolved and quantified. The extra control wells were used for contamination assessment in the whole confirmed workflow. C. Our optimized proteomics workflow profiled over 2,000 primary brain cells (diameter < 10 µm), beyond prior label free single-cell proteomics studies which focused on larger cultured cell lines or oocytes.

### Single-cell proteomics in the developing human brain

After strict data quality control, on average we quantified the abundance of approximately 800 distinct protein groups per cell in the brain (excluding erythroid cells), well above the background level estimated from empty microwells (**Fig. 1B**). This performance is particularly notable given the small size of developing human brain cells (approximately 10 μm in diameter) and is comparable with the typical gene coverage of 1,000–2,000 genes observed in single-cell RNA-seq experiments. In terms of scale, our profiling of whole proteomes from over 2,000 primary cells represents a significant advance over prior single-cell proteomics studies, that have primarily focused on cultured cell lines or relatively large cells such as oocytes ^13,20,21,25–31^ (**Fig. 1C** and **Table S1**). Notably, the GW13 sample exhibited a lower protein yield compared to later-stage samples, likely due to limited cellular diversity at early gestation. Importantly, we observed no significant difference in protein yield between cells isolated from the cortical plate and those from the germinal zone (**Fig. 1B**).

We performed clustering analysis in the UMAP space to group cells based on their proteomic profiles, identifying distinct clusters corresponding to erythroid and brain-associated cell populations (**Fig. 2A**). The erythroid cluster, uniquely marked by hemoglobin proteins alpha (HBA) and beta (HBB), separated into four major subclusters (**Fig. 2B**). Among these, fetal-specific hemoglobin zeta (HBZ) and adult-specific hemoglobin delta (HBD) clearly differentiated fetal and maternal erythroid cells, respectively, confirming the accuracy and resolution of our proteomic data (**Fig. 2C and D**). We particularly highlight a subgroup of fetal erythroid cells specifically expressing the ribosomal protein RPL6 (**Fig. 2E**). Prior studies established that ribosomal proteins are abundant in early erythroid progenitors, become strongly downregulated during reticulocyte maturation, and are entirely absent from mature erythrocytes due to the loss of organelles such as the nucleus, ribosomes, endoplasmic reticulum, and Golgi apparatus ^32^. Thus, the RPL6-positive subcluster identified here represents early-stage erythroid precursors. Additionally, our single-cell proteomic analysis demonstrated significantly lower protein diversity and reduced overall protein abundance in erythroid cells compared to brain-associated cells (**Fig. S1**), reflecting their specialized functional role dominated by hemoglobin and glycolytic enzymes, as well as their simplified cellular structure.

**Fig 2.**
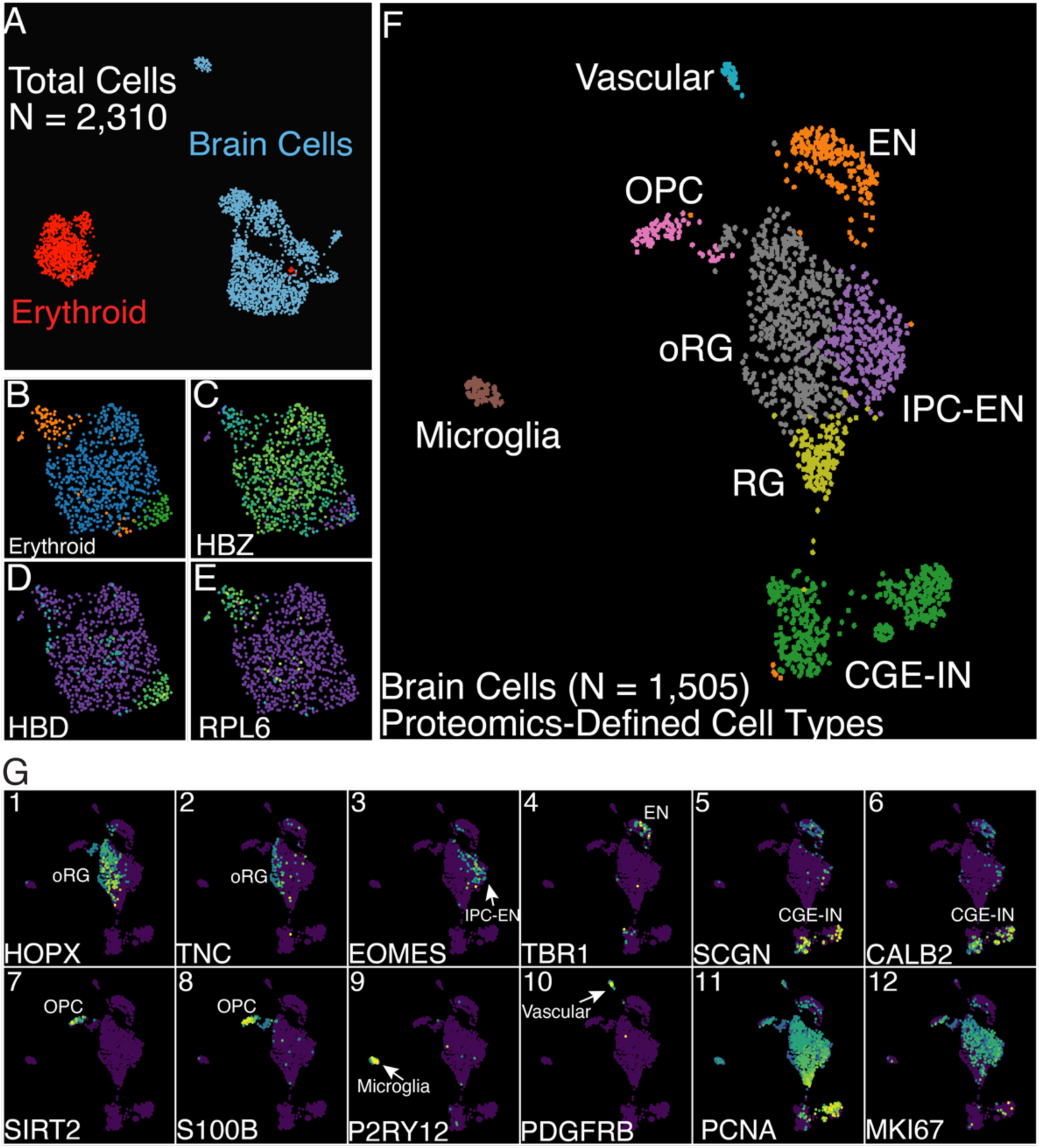
The Single-Cell Proteomic Atlas of the Developing Human Brain. A. A total of 2,310 single cells (Post-QC) were profiled, including erythroid (n = 805) and developing human brain cells (n = 1,505). B. High-resolution classification of the component of erythroid cells in fetal brain. C. Fetal-derived erythroid cells with high HBZ abundance, D. Maternal-derived erythroid cells marked by HBD protein. E. A highly transcriptionally active blood cells subcluster in fetal brain were identified with its high ribosomal protein RPL6 abundance. F. The proteomic single cell landscape of the developing human brain, including Radial Glial (RG) cells, outer Radial Glial (oRG) cells, Intermediate Progenitor/Excitatory Neuron (IPC-EN) cells, Excitatory Neurons (EN), GCE-Inhibitory Neurons (GCE-IN), Oligodendrocyte Progenitor Cells (OPC), Vascular and Microglia. G: The curated set of cell type markers used to define proteomic cell types in this study, including: HOPX and TNC for oRG, EOMES for IPC-EN, TBR1 for EN, SCGN and CALB2 for CGE-IN, S100B and SIRT2 for OPC, P2RY12 for Microglia, and PDGFRB for Vascular, PCNA and MKI67 for dividing cells.

We focused our subsequent analysis on 1,505 brain cells with resolved quantitative proteomes (**Fig. 2F**). Our clustering analysis identified main cell types in the developing brain, validated through established cell type markers (**Fig. 2G**): outer radial glia (oRG) cells were identified by HOPX and TNC (**Fig. 2G1-2**), whereas excitatory neuron (EN) progenitors and mature ENs were distinguished by EOMES and TBR1(**Fig. 2G3-4**), respectively. Additionally, we characterized caudal ganglionic eminence–derived interneurons (CGE-INs), specifically the SCGN-positive subtype, marked by SCGN and CALB2(**Fig. 2G5-6**). Oligodendrocyte progenitor cells (OPCs) were identified based on selective enrichment of S100B and SIRT2 proteins (**Fig. 2G7-8**), microglia were identified by P2RY12(**Fig. 2G9**), and vascular cells by PDGFRB (**Fig. 2G10**). Notably, cell-cycle proteins, PCNA and MKI67, markers respectively indicative of DNA replication and active proliferation, were predominantly observed in radial glia subtypes (RG/oRG), IPC-ENs and OPCs (**Fig. 2G11-12**). Interestingly, we discovered a subpopulation of CGE-INs exhibiting high PCNA levels but largely absent MKI67 (**Fig. 2G5-6,11-12**), suggesting these cells had recently exited the cell cycle into a G0 phase, yet retained residual PCNA expression. Further investigation revealed this unique CGE-IN subset was exclusively derived from the GW13 sample, that encompassed cells collected from the telencephalon, including the ganglionic eminence. Thus, these cells represent nascent CGE-INs from the CGE compartment. Collectively, our proteome-wide clustering robustly delineated distinct cell types; at the level of individual protein expression, marker-based validation confirmed the high quality of our dataset. Importantly, this dataset represents the first quantitative, single-cell–resolution proteomic profiling of the developing human brain.

### Comparative studies between single-cell RNA-seq and single-cell proteomics profiling

To uncover proteomic activity invisible to conventional single-cell RNA sequencing (scRNA-seq), we conducted comparative analyses using two independent scRNA-seq datasets. First, pairing our single-cell proteomics data, we generated single-cell transcriptomics data from the same prenatal brain samples. This dataset enabled direct, matched comparisons at the RNA and protein levels (**Fig. 3A-B)**. This paired RNA-seq data encompassed 31,639 cells and the recapitulated cell types are shown in **Fig. 3B** (see Materials and Methods). Second, to robustly account for inter-individual variability and to provide a broader biological context, we leveraged our recently published population-scale snRNA-seq reference atlas of the developing human brain^9^. This extensive dataset comprises single-nuclei transcriptomic profiles from a large cohort, spanning critical developmental stages from early gestation to adolescence, thereby establishing consensus gene-expression signatures for each major cell type (**Fig. 3C**). It is important to emphasize that single-cell RNA-seq has the capacity to profile substantially larger numbers of cells, which provides a robust foundation for defining high-confidence “expected” gene expression dynamics within each brain cell type. This extensive transcriptomic baseline, therefore, uniquely positions us to identify “unexpected” proteomic activity and molecular dynamics that are exclusively detectable through our single-cell proteomics approach, even when observed from a smaller population of cells.

**Fig 3.**
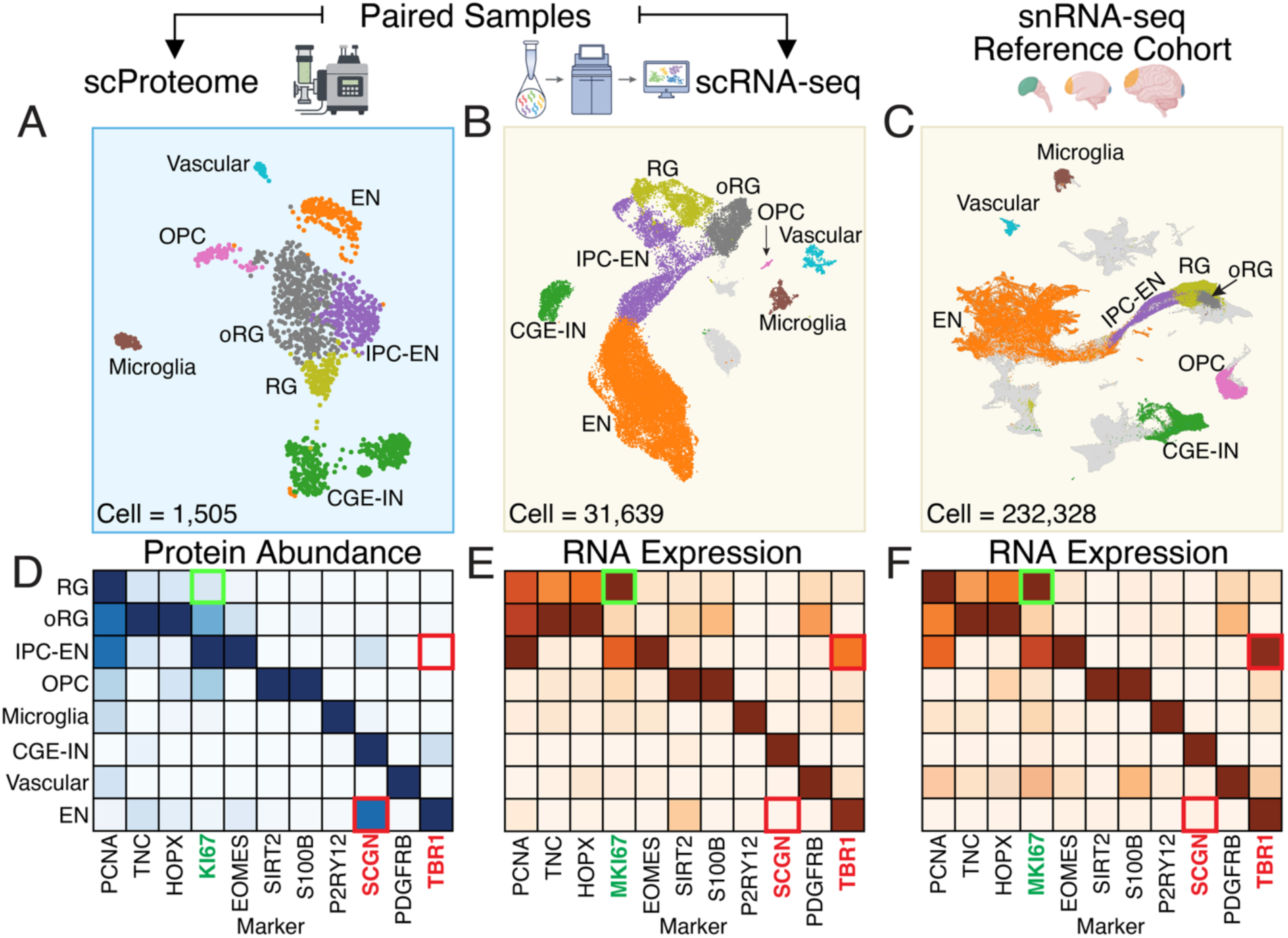
Comparative Analysis of Proteome and Transcriptomic Data Across Cell Types. A-B: Sample paired single-cell proteomics and RNA-seq data generated from the same developing human brains, C: Independent snRNA-seq validation cohort for comparison robustness. D-F: Canonical markers were selected to define and compare cell-type identities across both modalities. Notable discrepancies were highlighted with colored boxes. The gene MKI67 with differing turnover rates of the transcript and protein was highlighted in green boxed. TBR1 and SCGN, exhibited discrepancies in cell type specificity between mRNA and protein, were highlighted in red boxed.

We began by examining individual cell type markers. Many of these markers were originally characterized at the protein level in classical cell biology studies and were later adopted as cell type identifiers in single-cell RNA-seq analyses. As expected, their expression patterns were largely concordant across transcriptomic and proteomic datasets (**Fig. 3D-F**). However, notable discrepancies emerged that were validated experimentally (described below). For instance, in both the paired and reference RNA-seq datasets, the proliferation marker MKI67 exhibited the highest mRNA expression in radial glia (RG), followed by intermediate progenitor cells committed to excitatory neuron fate (IPC-EN) (**Fig. 3E and Fig. 3F**). In contrast, our single-cell proteomics data showed that the corresponding protein, KI67, was markedly reduced in RG, yet remained robustly expressed in IPC-EN (**Fig. 3D**). This discordance could be explained by the differing turnover rates of the transcript and protein: KI67 has a short protein half-life (∼1–1.5 hours)^33,34^ compared to a much longer mRNA half-life (∼5 hours)^35,36^, which would explain the strong MKI67 mRNA expression but much attenuated protein abundance in RG (**Fig. 3D vs Fig. 3E,F, green boxes**). Also notably, the low level KI67 protein in RG indicates that these cells have just exited the cell cycle, with residual PCNA protein still observable (**Fig. 3D PCNA in RG**) given PCNA stability through the early G0 stage. Together, these observations suggest that RG are entering a resting state and may be poised for subsequent differentiation. These developmental dynamics are only visible when a proteomic lens is integrated. In contrast, protein and mRNA observations were concordant in IPC-EN, where both data modalities revealed strong co-presence of KI67 and PCNA, indicating their active proliferation state (**Fig. 3D-F IPC-EN**).

Among the cell marker proteins, TBR1 and SCGN exhibited notable discrepancies between mRNA and protein levels (**Fig. 3D-F, red boxes**). TBR1 is a well-established marker of deep-layer cortical excitatory neurons (ENs)^37^, and as expected, our single-cell proteomics data faithfully captured specific TBR1 enrichment in ENs (**Fig. 4A3**). In contrast, single-cell transcriptomic data revealed strong TBR1 mRNA expression not only in ENs but also in intermediate progenitor cells committed to the excitatory lineage (IPC-EN) (**Fig. 4A1,** reference samples; **Fig. 4A2,** paired samples). We performed RNAscope and immunostaining to validate our observations: as expected, the TBR1 protein (by immunostaining, **Fig. 4A4**) and transcript (**Fig. 4A5**) are present in ENs in the cortical plate. In the germinal zone, IPC-EN cells (positive for the marker EOMES protein) displayed negative TBR1 protein (**Fig. 4A6**) but positive TBR1 mRNA expression (**Fig. 4A7**), faithfully recapitulating our observations from single cell protein profiling (**Fig. 4A3**). This mismatch, robust transcript levels in IPC-EN versus minimal detectable TBR1 protein, suggests a transcriptionally primed state in IPC-ENs, poised for rapid TBR1 protein synthesis upon differentiation into deep-layer neurons. Likewise, SCGN is a well-known marker for CGE-interneurons (CGE-IN)^38^, corresponding to its almost exclusive mRNA expression and its strongest protein abundance (**Fig.4B1-2**) in CGE-IN cells. However, in contrast to the almost complete absence of SCGN mRNA in non-CGE-IN cells, moderate or sometimes high SCGN protein abundance was detected in excitatory neurons (EN, **Fig.4B3**). We performed immunostaining for validation and, indeed, observed SCGN protein abundance in a subset of VGLUT1+ EN cells (**Fig. 4B4-6**), matching the observation from our single-cell proteomics profiling.

**Fig 4.**
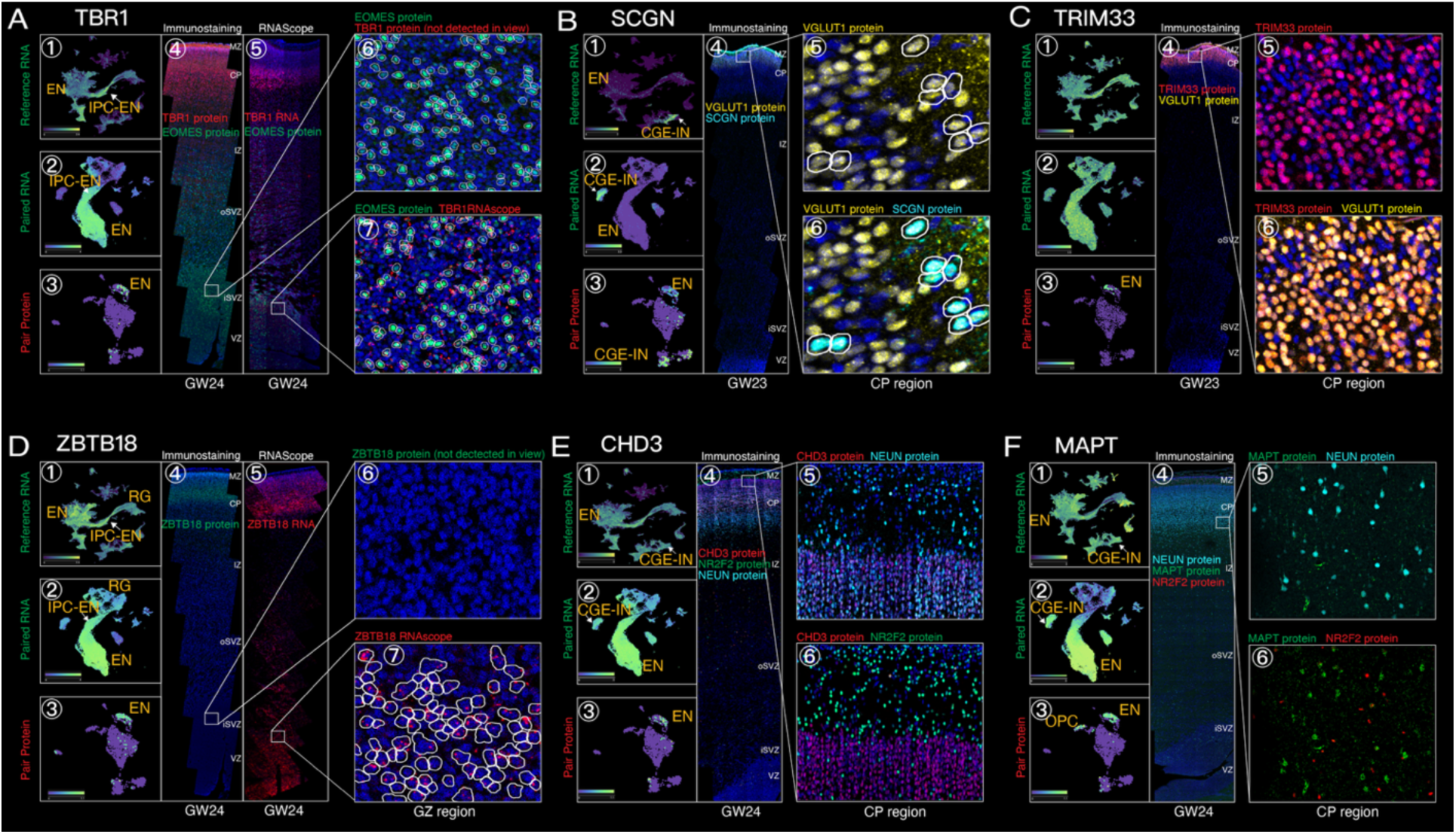
Immunostaining Validation of RNA-Protein Discordance in Cell-Type Specificity. A. **TBR1: Loss of cell-type specificity at the RNA level**. Protein expression is restricted to mature excitatory neurons (ENs) despite mRNA expression across intermediate progenitors (IPC-ENs) and EN cells. In zoom view (A6 and A7), IPC-EN cell borders are circled, TBR1 mRNA (detected via RNAscope) is present in IPC-EN cells, but no TBR1 protein product is detected, indicating a “primed-but-silent” pattern of gene regulation. B. **SCGN: Protein Expression Expands Beyond RNA-Specificity**. Mass Spectrometry and immunostaining results show the SCGN protein localizes in both EN cells (VGLUT1+) and CGE-IN cells(**B3-6**), while its mRNA restricted in CGE-INs(**B1-2**). C. **TRIM33: Novel Protein-Level Marker for EN Cells.** TRIM33 mRNA is broadly expressed across multiple cell types(**C1-2**), however, its protein is highly specific to EN cells(**C3**), validated by a co-localization of VGLUT1 and TRIM33 in EN cell(**C4-5**). D. **ZBTB18: Another novel EN-Cell Protein Marker.** Gene ZBTB18 shows general mRNA expression in RG, IPC-EN and EN cell(**D1-2**). However, its protein products are exclusively localized to mature EN cells(**D4**). Cells in GZ region express ZBTB18 mRNA are circled (**D7**), however, there is no ZBTB18 protein detected(**D6**). E. **CHD3: Autism-Associated Gene Activities in Specific Cell Type**. CHD3 mRNA is broadly expressed in both ENs and CGE-INs (**E1-2**), but Mass Spec and immunostaining result show its protein product is not detected in CGE-INs (**E3, E6**). F. **MAPT is absent in CGE-INs despite the presence of its transcript.** MAPT protein exist in EN cells and shows a contrast pattern with IN cells marker NR2F2.

The mRNA-protein discrepancies were even more pronounced when extending our analyses from known marker proteins to the global proteome (**Fig. S2)**. For example, TRIM33 mRNA expression is promiscuous across cell types in the developing cortex (**Fig. 4C1-2**), whereas protein expression from our single-cell profiling was strictly restricted to EN (**Fig. 4C3**). We performed immunostaining and validated that TRIM33 protein abundance was indeed confined to the cortical plate. Importantly, in the cortical plate, TRIM33 protein is co-present or co-absent with the EN marker, VGLUT1 (R= 0.93, P<0.0001), demonstrating its specific expression in ENs in the cortical plate, and replicating our observation from single cell proteomics profiling (**Fig. 4C4-6**).

We also performed validation on ZBTB18 whose mRNA was broad across cell types (**Fig.4 D1-2**) but whose protein was largely restricted to EN (**Fig. 4D3**). It was striking that while ZBTB18 mRNA was highly expressed across the cortical plate (**Fig.4D5**), ZBTB18 protein was confined to the upper layer (**Fig.4D4**), revealing mRNA-protein discordance. Moreover, in the germinal zone, moderate ZBTB18 mRNA expression was detected (**Fig. 4D6**), reflecting the scRNA-seq observation of attenuated but significant expression in radial glia and intermediate progenitor cells (**Fig. 4D7**). However, ZBTB18 protein was completely absent in the germinal zone (**Fig. 4D6**).

We particularly highlight CHD3, a gene implicated in both autism spectrum disorders (ASD) and Snijders Blok-Campeau syndrome^39,40^. CHD3 mRNA is broadly expressed, from radial glia to multiple neuronal subtypes (including ENs and INs), as well as in glial, microglial, and vascular cells (**Fig. 4E1-2**). This widespread transcriptomic profile has made it difficult to pinpoint the relevant cellular context for investigating CHD3’s pathogenic mechanisms. In contrast, our single-cell proteomics profiling revealed specific enrichment of CHD3 protein in ENs (**Fig. 4E3**). This protein abundance pattern was further validated by CHD3 immunostaining (**Fig. 4E4-6**), that showed restricted localization to the cortical plate and co-expression with the pan-neuron marker, NEUN. Furthermore, co-labeling using anti-NR2F2 showed non-overlap of CHD3 and NR2F2, demonstrating the absence of CHD3 in CGE-INs. The immunolocalization experiments faithfully captured our single-cell proteomics observations, establishing the EN-specific expression pattern of CHD3 at the protein level despite broad mRNA expression (**Fig 4E.1-2)**. Therefore, proteomic profiling pinpointed that the cellular basis of CHD3-associated conditions is related to ENs.

Similarly, our single cell data shed light on the cell-type specificity of MAPT, the well-known tau protein in Alzheimer’s disease, which is also implicated in neurodevelopmental disorders^41,42^. Single-cell RNA data revealed broad mRNA expression of MAPT across cell types (**Fig. 4F1-2**), particularly pronounced in ENs and INs, whereas our dataset indicates protein abundance is largely restricted to ENs and OPCs (**Fig. 4F3**). We performed immunostaining to verify whether high MAPT mRNA expression in IN cells, particularly CGE-INs, corresponds to high protein abundance in our single-cell proteomics data. Replicating our proteomics profiling observation, MAPT protein abundance was specific to cortical neurons (NEUN+, **Fig. 4F5**) but was absent from CGE-INs (marked by NR2F2, **Fig. 4F6**), in contrast to strong MAPT mRNA expression in CGE-INs in both reference (**Fig. 4F1**) and paired RNA-seq datasets (**Fig. 4F2**). These experiments validated our single-cell proteomic observation, providing a further example of mRNA and protein discordance.

Taken together, our experimental validations confirm the high quality of the single-cell proteomics data. More importantly, the observed discrepancies between mRNA and protein abundance underscore the critical need to study disease-associated genes at the protein level, as reliance on transcriptomic data alone may lead to misinterpretation of their cellular specificity and functional relevance.

### Neurodevelopmental genes display discordant mRNA expression and protein abundance

The discordance between mRNA and protein expression led us to evaluate how well mRNA levels predict protein abundance in each cell type. For this analysis we integrated matched single-cell proteomic and transcriptomic datasets. Protein quantification in our single cell analysis relied chiefly on DIA-NN^24^, which offers unbiased cell-to-cell measurement of individual proteins. However, between-protein comparisons can be skewed by potential differences in peptide detectability. To correct for this, we also applied the iBAQ method ^43^, which normalizes for the number of theoretically observable peptides and thus approximates absolute protein abundance. Despite their different algorithms and objectives, DIA-NN and iBAQ produced highly concordant results in our data (median Spearman ρ = 0.902; mean = 0.907; P < 0.001). Because DIA-NN is specifically optimized for single-cell analysis, we report these results in the main text, with parallel iBAQ validations provided in Methods. Conclusions were consistent across both approaches.

We observed that proteome and transcriptome correlations were strongly cell-type dependent: excitatory neurons (ENs) showed the highest mRNA–protein concordance, whereas interneurons (INs) exhibited the lowest (**Fig. 5A**). Even in ENs, however, the average correlation was modest (Spearman’s ρ = 0.36, P < 1e-3). These findings align with bulk-tissue studies^3^ and underscore the limits of transcriptomics for inferring protein expression, reinforcing the need for direct single-cell proteomic profiling. Within each cell type, we further applied regression analysis to identify extreme mRNA–protein outliers (P ≤ 0.05, see Materials and Methods): one group of genes displayed low RNA but high protein levels (Group A), while the reciprocal group showed high RNA but low protein levels (Group B). Representative outlier genes in ENs are illustrated in **Fig. 5B**. Group A and B genes were identified in each cell type in the developing brain (**Fig. 5C1).** We asked what genes are more likely to have highly discordant mRNA expression and protein abundance. Group-A genes, characterized by low mRNA expression but high protein abundance, exhibited strong functional enrichment for key regulatory pathways including stem cell differentiation, mRNA splicing, and anti-apoptotic mechanisms (**Fig. 5D**). Analysis using the Mouse Genome Informatics database further revealed significant enrichment of phenotypes such as abnormal cell-cycle progression and disrupted cortical stratification among mutants of these genes (**Fig. 5D**). Such enrichment suggests that the observed discordance between low transcript levels and high protein abundance might represent an adaptive regulatory mechanism. Specifically, maintaining abundant and stable protein pools from limited mRNA could ensure rapid and reliable cellular responses under fluctuating transcriptional conditions. This regulatory paradigm is exemplified by splicing factors, which frequently autoregulate their own transcripts via alternative splicing coupled to nonsense-mediated decay, thus reducing mRNA abundance while maintaining robust protein reservoirs^44^.

**Fig 5.**
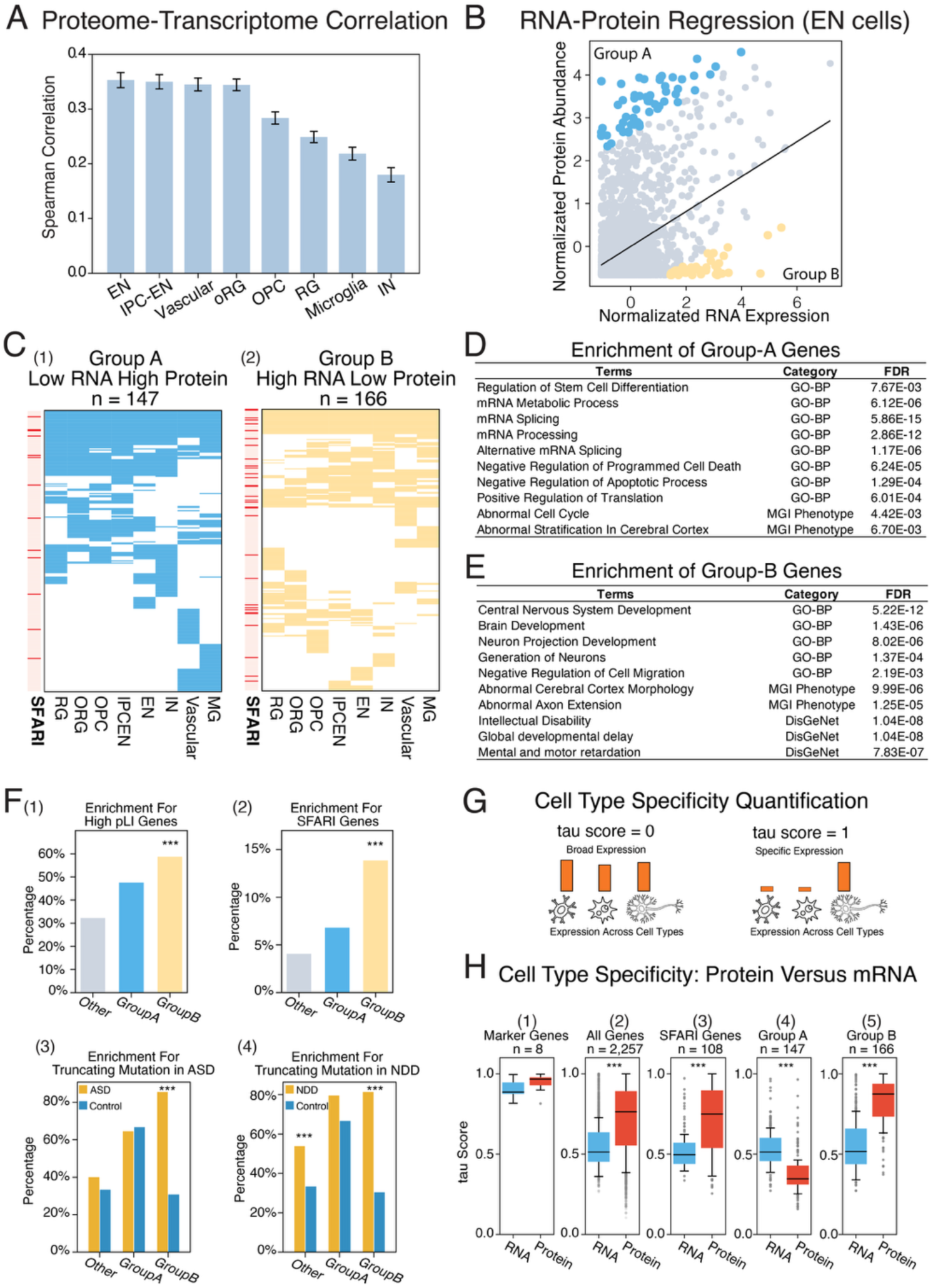
Integrated Sample-Paired Protein-RNA Profiling Uncovers Widespread Discordance in Gene Expression and Cell Type Specificity. A. Proteome-transcript comparison shows global low correlation between RNA and protein across different cell types. B. Illustration of the linear regression used to identify extreme outlier gene groups in EN cells (p < 0.05). Group A genes exhibit low RNA but relatively high protein abundance, whereas Group B genes show high mRNA expression but unexpectedly low protein abundance. C. Pattern of all outlier group A genes (n = 147) and outlier group B genes (n = 166) across all cell types. SFARI bar indicates the SFAREI genes distribution in each group. D. Group-A genes show strong functional enrichment in mRNA splicing, stem cell differentiation, and anti-apoptotic mechanisms. E. Group-B genes show significant enrichment for critical neurodevelopmental processes. F. Compared to other inlier protein, Group B contains more high-pLI genes (1) and enriches in more SFARI genes (2). And moreover, Group B genes show higher truncating mutation enrichment in both the ASD and NDD cohort than the control cohort (3-4). ASD: Autism, NDD: Neurodevelopmental Diseases. G. Illustration of the tau score, an index of cell-type specificity. Genes with broad expression will be assigned a tau score close to 0, whereas cell-type-specific genes have a tau score close to 1. H. Comparison of cell-type specificity at the protein and RNA levels using tau score across five gene sets: (1) marker genes (n = 8), (2) all quantified genes in paired sample (n = 2,257), (3) All quantified SFARI genes in this study (n = 108), (4) 147 Group-A gene with low RNA but high protein pattern and (5) 166 Group-B gene with high RNA but low protein pattern. Higher tau scores indicate greater cell-type specificity.

In contrast, Group B genes exhibited high mRNA expression but unexpectedly low protein abundance (**Fig. 5C2**), suggesting that extensive post-transcriptional or translational regulation is required to fine-tune and restrict protein levels to meet precise biological needs. Several lines of evidence strongly argue against a technical shortfall in our proteomics profiling that may have failed to capture these proteins. First, our analyses were performed at a pseudobulk cell-type level, meaning that consistently low protein abundance across numerous independently sampled cells of the same type precludes stochastic sampling artifacts. Second, given that our mass spectrometry workflow achieves true single-cell sensitivity, proteins remaining undetected or scarce indeed reflect genuine biological scarcity rather than technical limitations, particularly striking when compared to their abundant mRNA counterparts. Third, in situ immunostaining experiments independently validated this pattern, confirming widespread mRNA expression but restricted protein localization (**Fig. 4**). Supporting this further, we identified the 166 proteins displaying this discordant trend consistently across multiple cell types (**Fig. 5C2**), and these were strongly enriched for proteins with known rapid degradation kinetics and short half-lives^45^ (P = 0.0035). Collectively, these observations affirm that our analysis accurately captured a biologically meaningful class of proteins whose rapid turnover overrides abundant transcriptional input.

Functionally, Group-B genes showed significant enrichment for critical developmental processes such as neuron generation, neuronal migration, and broader neurodevelopmental programs (**Fig. 5E**). Mutant mouse models for these genes frequently exhibit aberrant cortical morphology and disrupted axonal outgrowth, underscoring their roles in fundamental aspects of neurodevelopment (**Fig. 5E**). We reasoned that the attenuated protein abundance relative to their mRNA expression likely suggests dosage sensitivity of the Group-B proteins. We thus examined the probability of loss-of-function intolerance (pLI) scores^46^, a metric reflecting sensitivity to dosage alterations and intolerance to truncating genetic mutations. Indeed, Group-B proteins displayed significantly higher pLI scores compared to Group-A proteins and other inlier proteins (with concordant mRNA and protein levels) (**Fig. 5F1**). Thus, elevated pLI scores confirmed that Group-B proteins are dosage-sensitive, reinforcing our interpretation that the discrepancy between abundant transcripts and scarce proteins represents a tightly regulated biological mechanism rather than experimental noise. To determine clinical associations, we first examined high-confidence autism spectrum disorder (ASD)-associated genes from the SFARI database^47^ (confidence levels 1 and 2) and detected a significant enrichment (**Fig. 5F2**) among Group-B protein relative to Group-A or to other inlier proteins. Instead of using this curated ASD gene set, we further examined agnostically identified de novo mutations from whole-genome sequencing for individuals with ASD as well as their unaffected siblings from 37,368 families in a recent study^48^. We examined proteins with de novo mutations across Group A, Group B, and the inlier set (the *Other* group, **Fig. 5F3**), calculating the fraction of proteins harboring truncating mutations (stop-gain/loss, splicing, or frameshift) in each group among those with detectable de novo mutations. We then compared these fractions between ASD cases and unaffected siblings to assess mutational enrichment. Notably, only Group B proteins showed a distinctive enrichment for truncating mutations in ASD cases (**Fig. 5F3**), further supporting their association with autism risk. Lastly, the same comparison was performed on de novo mutations identified from large-scale probands with general neurodevelopmental disorders^48^. Both Group-B and the inlier set displayed greater enrichment for truncating mutations in NDD probands than unaffected controls (**Fig. 5F4**). Taken together, proteins in Group B are strongly implicated in neurodevelopmental disorders, particularly ASD. The combination of high mRNA expression but low protein abundance suggests that precise post-transcriptional and/or translational regulation is critical for maintaining the physiologically appropriate protein levels of these ASD-associated genes. This underscores their dose sensitivity, as highlighted in **Fig. 5F1**, and aligns with the well-established role of RNA-binding proteins and post-transcriptional control in neuropsychiatric disorders^49^. Given this, relying solely on high mRNA expression as an indicator of molecular activity would mask the underlying biology of these genes—specifically, their rapid protein degradation or tightly restricted translation—potentially leading to misinterpretation of their functional roles. Notably, as shown in Fig. 5C2, mRNA–protein discordance was often cell-type-specific, suggesting a mechanism by which protein production is selectively dampened to establish or reinforce cell-type identity, as we further detail below.

### Protein abundance is more cell-type-specific than mRNA expression

In addition to analyzing mRNA–protein discordance within individual cell types, we further compared cell-type specificity at both mRNA and protein levels. We employed the widely utilized tau score^50^ to quantify cell-type-specific gene expression or protein abundance across multiple cell types (based on mean expression at the pseudobulk cell-type level). A tau score approaching 1 indicates highly cell-type-specific expression, whereas a score near 0 implies ubiquitous expression across cell types (**Fig. 5G**). We initially examined established marker genes/proteins for major brain cell types (**Fig. 5H1**) and confirmed strong cell-type specificity, with tau scores above 0.9 for both mRNA and protein modalities. However, upon extending this analysis to all detected proteins in our single-cell proteomic dataset, we observed markedly greater cell-type specificity at the protein level compared to mRNA expression derived from paired tissue samples (**Fig. 5H2**) and from an independent reference cohort (**Fig. S4**). This difference was validated by immunostaining experiments, where mRNA showed broader, more promiscuous expression patterns, whereas protein expression remained confined to distinct cell types (**Fig. 4**). Notably, autism spectrum disorder (ASD)-associated proteins mirrored this overall proteomic trend, exhibiting significantly higher cell-type specificity at the protein level than at the mRNA level (**Fig. 5H3**).

We highlight two examples, PHIP and NOVA2, to illustrate this key point. PHIP is strongly associated with neurodevelopmental disorders^39,51^, including Chung-Jansen syndrome^52^, yet its etiological mechanisms have remained unclear. Our single-cell proteomics data pinpointed PHIP’s primary functional localization in radial glia (RG), outer radial glia (oRG), and intermediate progenitor cells transitioning to excitatory neurons (IPC-EN) (**Fig. S5A**). In stark contrast, PHIP exhibited widespread, non-specific mRNA expression patterns in both our paired samples (**Fig. S5B**) and reference cohorts (**Fig. S5C**). Similarly, NOVA2, another ASD-associated gene^53^, displayed clear enrichment in excitatory neurons (EN) and oligodendrocyte precursor cells (OPC) at the protein level (**Fig. S5D**), despite broad, cell-type-agnostic mRNA expression profiles (**Fig. S5E** and **S5F**).

Despite the general trend, the two groups of outlier proteins (Group A and Group B) exhibited opposite behaviors. Group A genes were characterized by low mRNA expression but high protein abundance across cell types (**Fig. 5C1**). This elevated protein abundance dampened their cell-type specificity, as reflected by substantially lower tau scores at the protein level compared to the mRNA level (**Fig. 5H4** vs. **Fig. 5H2**). In contrast, Group B genes, defined by high mRNA expression but low protein abundance, displayed significantly higher cell-type specificity at the protein level than the global proteome background (median tau scores: 0.86 for Group B vs. 0.76 for background, P < 0.001, Wilcoxon rank-sum test). This disproportionate reduction in protein levels sharpens their cell-type specificity without altering specificity at the transcript level (**Fig.5H5** vs **Fig.5H2**). Collectively, these findings highlight how protein-level analysis reveals critical mechanistic insights—such as the contribution of translational control to cell identity—that remain hidden when relying on transcriptomic data alone. Our results emphasize the necessity of integrating single-cell proteomics into neurobiological research to achieve a more accurate and complete understanding of disease-associated genes and mechanisms.

### Lineage reconstruction – Identifying a protein module in autism spectrum disorders

At the protein level, we reconstructed the cell developmental trajectory from radial glia (RG) to excitatory neuron (EN) differentiation through the intermediate progenitor state (IPC-EN, **Fig. 6A**). Developmental dynamics across pseudotime recapitulated the characteristic transient up-regulation of EOMES in IPC-EN and later activation of EN markers TBR1 and ZBTB18 (**Fig. 6B**). We implemented Weighted Gene Co-expression Network Analysis (WGCNA)^54^ across developmental pseudotime, and the time-course analysis identified six co-expressed protein modules, displaying time-dependent activities along the RG→EN developmental trajectory (**Fig. 6C**). For example, Module 1 encompassed proteins active in RG with declining protein abundance as development progresses. Module 4 displayed activation beginning at IPC-EN, whereas Module 5 showed activation from the EN stage onward.

**Fig 6.**
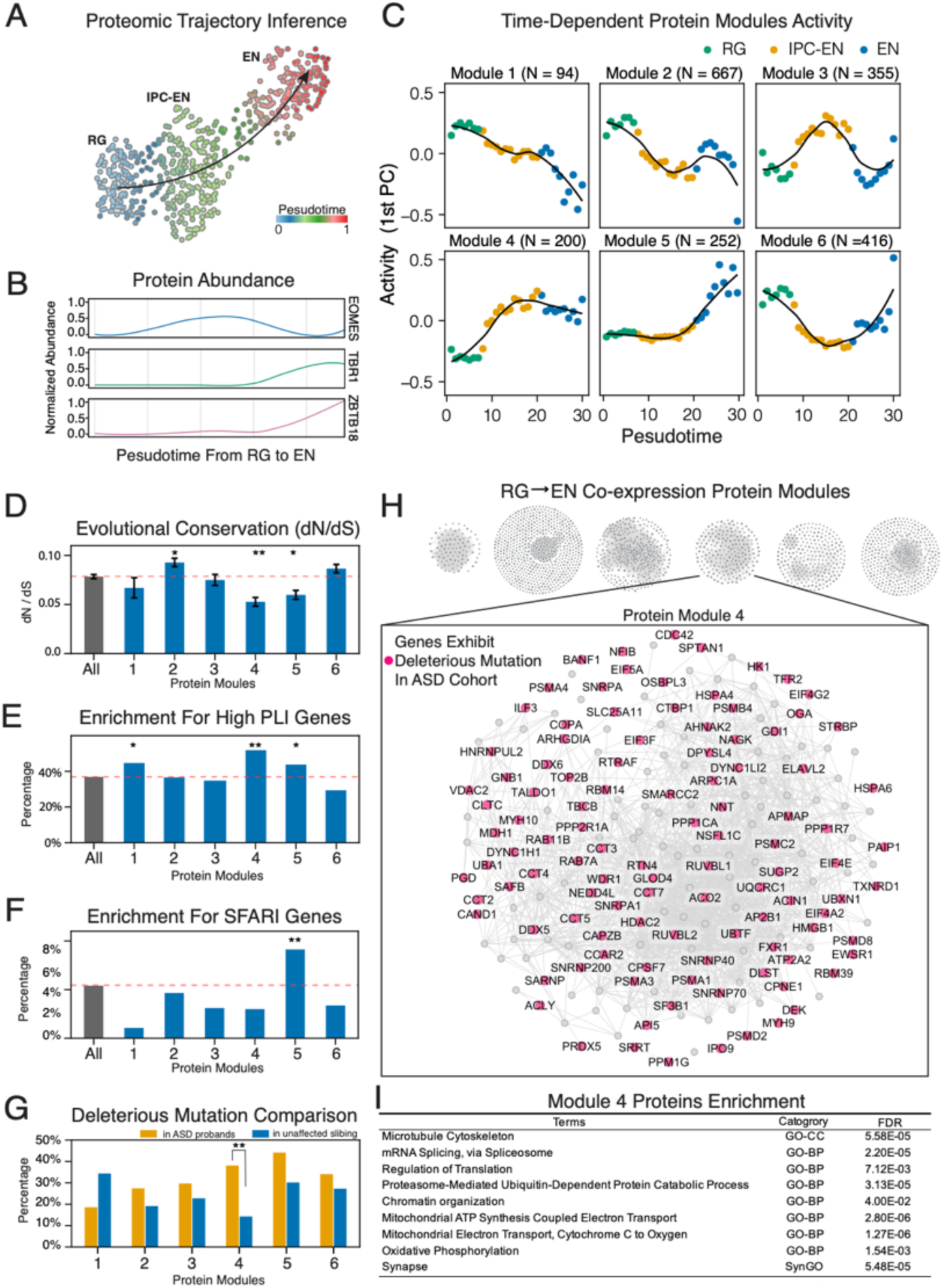
Protein Dynamics in Human Neurodevelopment. A. Proteomic trajectory inference of EN development. The trajectory traces human EN cell development from RG to IPC-EN to mature EN cell, using 532 single-cell protein profiles (129 RG cells, 232 IPC-EN cells and 171 EN cells). B. Proteomic trajectory recapitulates the characteristic transient up-regulation of EOMES in IPC-EN and later activation of EN markers TBR1 and ZBTB18. C. WGCNA reveals 6 protein modules driving developmental stages, module 4 activity represents proteins active in the IPC-EN/EN junction stage, showing high activity at the transition from IPC/EN to neurons, and proteins in module 5 show a rapid activity during the EN maturation. D. Evolutionary rates (dN/dS) analysis reveals a high conservation in module 4 and 5, compared to Mus Musculus, underscores the functional necessity of proteins in module 4 and 5 for basic neurodevelopment. E. Evolutionary constraint of genes in module 4 and 5 is further supported by their higher intolerance to protein-truncating mutations than other genes. F. Genes in module 5 contains more genes associated with autism spectrum disorders (ASD), P < 0.01. G. Genes in module 4 carry more deleterious missense mutations in ASD probands relative to unaffected sibling controls (P < 0.01). H. The protein co-expression network in human neurodevelopment process. The module 4 is zoomed in and genes with deleterious missense mutations in ASD cohort were highlighted in red. I. The GO terms enrichment for the proteins in module 4.

For proteins in each module, we identified their one-to-one mouse orthologs and computed their respective evolutionary rates, ω (the dN/dS ratio; see Materials and Methods), since the mouse–human split. Strikingly, we observed that Modules 4 and 5 displayed substantially reduced ω, highlighting their strong evolutionary conservation (**Fig. 6D**). This evolutionary constraint is further supported by human population genomic data: both modules were highly enriched for proteins with extreme pLI scores (**Fig. 6E**)^46^, indicating intolerance to protein-truncating mutations. When examining genes associated with autism spectrum disorders (ASD) curated by SFARI (confidence scores above 2), Module 5 displayed unique enrichment (**Fig. 6F**). This is consistent with the known SFARI genes enriched for synaptic functions, consistent with the functional profile of Module 5 which becomes active during EN emergence (**Fig. 5C**).

To achieve an agnostic genomic scan, we leveraged the largest dataset to date for de novo mutations in ASD probands and their unaffected siblings^48^. Although truncating mutations were at a low level in Module 4, likely reflecting strong dosage sensitivity indicated by its high pLI scores, analysis of deleterious de novo missense mutations revealed significant enrichment in ASD probands compared to unaffected sibling controls (**Fig. 6G**). This excess of damaging mutations in ASD cases underscores a robust association between Module 4 and ASD vulnerability.

Examining the specific proteins in Module 4 revealed many known ASD-associated genes (**Fig. 6H**), such as RUVBL2, SMARCC2, NFIB, and CTBP1^39,55–57^. The activity of Module 4 initiates at the IPC-EN transition, marking a key developmental window of vulnerability to ASD. Although Module 4 proteins are enriched for multiple GO terms (**Fig. 6I**), they converge on a single, interlocking workflow that transforms a mitotically active, metabolically glycolytic progenitor into a post-mitotic neuron capable of migration, arborization, and synaptic signaling. Briefly, the transition from intermediate progenitor cells (IPCs) to excitatory neurons (ENs) involves a tightly coordinated, multi-layered cellular transformation. Structurally, neurons reorganize their cytoskeleton through dynein motors (DYNC1H1, DYNC1LI2), LIS1 (PAFAH1B1), Arp2/3 complex subunits (ARPC1A, ARPC4), actomyosin motors (MYH9, MYH10), and the chaperonin ring (CCT2–8, CCT6A), enabling migration and dendritic arborization. Simultaneously, a comprehensive shift in RNA processing occurs via spliceosome components (SF3B1, SNRPD2, SRSF2, U2AF1, SNRNP200), RNA helicases (DDX5, DDX6), and translation initiation factors (EIF1, EIF4E, EIF4G2), creating neuron-specific isoforms essential for synaptogenesis^58^. Enhanced proteostasis mechanisms, involving proteasome subunits (PSMA1–7, PSMC2, PSMD2/8), ubiquitin modifiers (NEDD8, NEDD4L, USP5), and chaperones (HSPA4, CCT5), ensure precise regulation of protein synthesis and degradation^59^.

Chromatin remodeling proteins (ACTL6A, SMARCC2, histone H1 isoforms H1-1, H1-4, DEK, HCFC1, CTBP1, PPP2R1A, CCAR2) reset transcriptional programs to enforce neuronal identity. This elaborate transformation is metabolically supported by a shift from glycolysis to oxidative phosphorylation, driven by mitochondrial proteins (MT-CO2, COX4I1, COX6C, COX7A2, NDUFS8), tricarboxylic acid cycle enzymes (IDH1, ACO2), and metabolic regulators (NNT, VDAC2/3)^60^. Finally, specialized trafficking proteins (CLTC, CLTA, AP2B1, RAB7A, RAB11B, GDI1, PGRMC1, OSBPL3) establish vesicle dynamics and synaptic functionality. Together, these integrated processes outline a seamless developmental progression from IPCs to mature ENs, emphasizing sustained functional roles in neuronal connectivity, metabolic stability, and synaptic maintenance.

Taken together, our WGCNA analysis at the proteomic level revealed Module 4 and 5 in ASD. While the identification of Module 5 was consistent with our knowledge of the role of EN in this disease, the identification of Module 4 was significant, revealing the IPC-EN transition as a developmental bottleneck in ASD pathogenesis, consistent with findings from our prior single-cell epigenomic study^9^.

## Discussion

Single-cell proteomics has long been thought of as the next frontier for charting cellular complexity in human tissues, yet, beyond cultured cell lines and oversized primary cells such as oocytes or hepatocytes, technical barriers have stalled its broad in vivo adoption. Here, we overcome those barriers with an optimized SCP workflow that reliably quantifies thousands of proteins in small physiologic cells, including prenatal neurons only 4–10μm in diameter. This advance transforms single-cell proteomics from a proof-of-concept technology into a practical platform for large-scale proteomic mapping of complex human tissues.

Our high-sensitivity single-cell proteomics map reveals the protein landscape of individual cells in the developing human cerebral cortex. A direct comparison with matched single-cell transcriptomes shows that RNA levels are an imperfect proxy for protein abundance: even in excitatory neurons, where concordance is highest, the mRNA-to-protein correlation remained below 0.4 (Spearman’s ρ = 0.36, **Fig. 5A**). This pervasive discordance underscores why protein-level measurements are essential for decoding developmental mechanisms and disease risk.

Two outlier gene classes illustrate how post-transcriptional control shapes cell identity. The first group displays low mRNA but high protein levels and is enriched for spliceosomal components (**Fig. 5D**). Because splicing factors must be present in large excess, cells appear to achieve this by maintaining a modest basal pool of transcripts while keeping protein synthesis high, an economical strategy that would be invisible in RNA-only studies. The second group shows the opposite pattern: high RNA but low protein. These genes cluster into neurodevelopmental functions that are strikingly enriched for autism-spectrum-disorder (ASD) loci. The underlying mechanisms are diverse. For the deep-layer marker TBR1, we observe strong transcription in intermediate progenitor cells transitioning to excitatory neurons (**Fig. 4A1-2 and Fig. 4A7**), yet TBR1 protein was undetectable (**Fig. 4A6**). Such “primed-but-silent” expression poises the cells for a rapid post-mitotic switch to neuronal identity. Other members of this group undergo rapid protein turnover across all lineages (**Fig. 5E**), indicating active degradation independent of cellular context.

Quantitatively, protein abundance proves to be far more cell-type-specific than mRNA expression (**Fig. 5H**). Transcriptional bursts^61^ make RNA levels stochastic and broadly distributed, whereas protein output resolves into clearer ON/OFF states once translation and degradation are taken into account. This distinction carries clinical significance. For many genes associated with neurodevelopmental diseases, the level of protein expression unmasks lineage-restricted vulnerabilities that may be masked by promiscuous RNA expression. This is best exemplified by several ASD-associated proteins (e.g., PHIP, NOVA and CHD3, **Fig. 4E** and **Fig. S4**) that show strong cell-type bias at the protein level despite ubiquitous transcripts. These findings place post-transcriptional and translational regulation at the forefront of human corticogenesis and neurodevelopmental pathology, providing strong support for the long-suspected contribution of post-transcriptional and translational regulation to neurodevelopmental disorders ^62–64^. These findings caution against interpreting disease biology from transcriptomics alone and illustrate how single-cell proteomics can pinpoint the precise cell types where therapeutic intervention might be most effective.

Leveraging our single-cell proteomics data, we reconstructed the developmental trajectory from radial glia to excitatory neurons, enabling a proteome-resolved view of cortical neurogenesis (**Fig. 6A**). This approach revealed a co-regulated protein module that is highly enriched for genes disproportionately affected by de novo mutations in individuals with autism spectrum disorders (ASD). Remarkably, the coordinated activity of this module emerges specifically during the transition from intermediate progenitors to post-mitotic excitatory neurons, a developmental window exactly identified in our GWAS-based analysis as a peak period of autism risk^9^. The convergence of findings from both common and de novo variant studies strongly supports a shared pathophysiological mechanism in ASD centered on the progenitor-to-newborn neuron transition.

Closer analysis of Module 4 revealed a highly orchestrated molecular program that initiates at the IPC-to-EN transition—precisely when lineage commitment and early circuit assembly begins. Rather than a random assortment of ASD-associated genes, Module 4 proteins form a coherent developmental engine that drives the transformation of dividing progenitors into functionally mature excitatory neurons. This transformation spans multiple biological layers: structural reorganization via cytoskeletal remodeling complexes, transcriptomic rewiring through chromatin remodelers and splicing machinery, metabolic shifts favoring oxidative phosphorylation, and the establishment of synaptic vesicle trafficking systems. The convergence of these processes not only ensures successful neuronal maturation, but also highlights how disruptions in any component, many of which are known ASD risk genes, can derail this tightly timed developmental progression. This modular framework thus provides mechanistic insight into how diverse genetic mutations may converge on a shared vulnerability window during human corticogenesis. We particularly note the remarkable evolutionary conservation in Module 4 (**Fig. 6D**), suggesting that the coordinated molecular program driving the progenitor-to-neuron transition is fundamental to vertebrate brain development.

Taken together, this study delivers the first proteome-wide map of individual cells in the developing human cortex, showcasing the unique power of single-cell proteomics to reveal biology hidden from RNA-based approaches. This study establishes that single-cell proteomics is no longer aspirational but is a practical tool that can reshape our understanding of human brain development and disease. As the technology continues to mature, gaining depth, speed, and accessibility, it promises to propel the field forward, enabling precise dissection of cellular states, pathways, and therapeutic targets that have remained out of reach with genomics and transcriptomics alone.

## Materials and Methods

### Samples Collection and Data Generation

We obtained three fresh, prenatal human brain tissue at gestational weeks (GW) 13, 15, and 19 from Zuckerberg San Francisco General Hospital (ZSFGH). Acquisition of human tissue samples was approved by the UCSF Human Gamete, Embryo and Stem Cell Research Committee (10-05113). All experiments were performed in accordance with protocol guidelines. Informed consent was obtained before sample collection and use for this study. Cortical tissue samples from GW15 and GW19 were micro-dissected into germinal zone (GZ) and cortical plate (CP). For sample at the early developmental stage of GW13, we used the whole frontal telencephalon, including ganglionic eminence. All samples were immediately placed in Hibernate-E medium (Invitrogen) prior to further processing. Label-free single-cell mass spectrometry and single-cell RNA sequencing were used for proteomic and transcriptomic profiling. Data quality control and analysis were conducted using Seurat and custom scripts, with trajectory analysis fitted using Scanpy.

### Immunohistochemistry and RNAscope

Primary human cortical tissue was collected and processed as approved by UCSF Gamete, Embryo and Stem Cell Research Committee (GESCR). All samples were de-identified with no information on sex. Tissue was collected with patient consent for research following strict legal, institutional, and ethical regulations. Primary cortical tissue samples from second trimester were fixed using paraformaldehyde overnight at 4°C, washed with 1X PBS, cryopreserved using 30% sucrose and sectioned onto FisherbrandTM SuperfrostTM (16um). Immunostaining was performed following the protocol described in prior publication^65^. The following antibodies were used: TBR1 (MilliporeSigma, AB2261, 1:300), SCGN (Arigo Biolaboratries, ARG11060, 1:300), TRIM33 (Millipore, MABE432, 1:200), ZBTB18 (Atlas Antibodies, CHM05N496, 1:300), CHD3 (Abcam, ab109195,1:300), MAPT (Cell Signaling, 66850S, 1:300). EOMES (R&D Systems, AF6166, 1:250), NEUN (Millipore, ABN90,1:500), NR2F2 (Novus Biologicals, PP-H7147-00, 1:300), VGLUT1 (Synaptic Systems, 135 302,1:300).

The RNAscope Multiplex Fluorescent Reagent Kit v2 were purchased from ACD Advanced Cell Diagnostics. RNAscope was conducted for TBR1 and ZBTB18 according to the manufacturer’s recommendation. Probes for TBR1 (Hs-TBR1; Catalogue No., targeting region) and ZBTB18 (Hs-ZBTB18; Catalogue No., targeting region) were purchased from ACD biosciences. Opal 570 (Akoya Biosciences) was used as a fluorophore. After sections were stained, confocal images were acquired with a Leica TCS SP5 X. Acquired images were processed using ImageJ^66^.

## Statistical Analysis

GO enrichment analysis was performed using Enrichr^67^ to query the Gene Ontology (Biological Process), MGI phenotype, and DisGeNet databases. Statistical significance of enrichment was determined using a hypergeometric test, with p-values adjusted for multiple testing using the Benjamini-Hochberg method. Enrichment for short-live protein, enrichment for SFARI genes^68^, enrichment for genes with high probability of being loss-of-function intolerant (pLI)^46^, and enrichment for genes with truncating mutations was assessed using functions implemented in custom script.

